# Lung Epithelial TLR7 is a Factor in Viral Replication and Pathogenesis

**DOI:** 10.64898/2026.11.03.764012

**Authors:** Madison J. Granoski, Ryley D. McClelland, Leanne M. Bilawchuk, Stephanie Hindle, Joaquin Lopez-Orozco, Quinten Kieser, Lionel D. Jensen, Cameron D. Griffiths, Farah Elawar, Olivier Julien, Deanna M. Santer, Carola G. Vinuesa, Tom C. Hobman, Craig N. Jenne, David J. Marchant

## Abstract

Respiratory virus infections are the leading causes of morbidity and mortality of infants and the elderly worldwide. Currently it is not known how Respiratory Syncytial Virus (RSV) vaccination of infants can cause enhancement of RSV disease, highlighting gaps in our understanding about how viruses like RSV interact with the immune system. Here we address this and show that ERK MAP kinase signaling from Toll-like Receptor-7 (TLR7) detection of RSV and influenza, but not human parainfluenza virus type 3 (HPIV3), infections in bronchial epithelial cells (BECs) drives replication of the viruses. Virus infection is detected by pattern recognition receptors like TLR7 that bind to single stranded RNA (ssRNA) viral replication intermediates in late endosomes. Here we report that *TLR7* is expressed in BECs and detects RSV ssRNA, causing ERK MAP kinase signaling. RSV and influenza use this detection and signaling in BECs to drive its replication and infection, a mechanism that counters TLR7s role in antiviral defence. Experimental blockade of TLR7 binding or downstream signalling through ERK MAP kinase decreased RSV and influenza replication, but had no effect on HPIV3. RSV viral loads in mouse and human females were higher compared to males, consistent with higher sex-dependent expression of the *TLR7* gene from two copies of the X chromosome in females. In a TLR7 gain of function mouse model there was a significant enhancement of RSV replication and RSV eosinophilic and Th2 disease markers like interleukin-5 (IL-5). In summary, TLR7 is expressed in human lung BECs in the lungs where it acts as a factor in RSV and influenza replication and disease pathogenesis. We propose that this is a major factor in determining the sex-disparity of viral disease.

---

Respiratory Syncytial Virus (RSV) is the leading cause of hospitalization of infants worldwide. RSV vaccine trials in the 1960s led to the hospitalization of up to 80% of the vaccinees and the deaths of two young boys^1^, and more recently, an mRNA vaccine trial caused enhanced RSV disease in infant vaccinees^2^. It is still not understood how the host response to vaccination can enhance RSV disease, highlighting a significant gap in our understanding of the host response to RSV infection.

RSV uses receptors and their signaling to mediate its entry into host cells^3^ and here we looked for receptors that are required for RSV replication after entry. We first identified the primary receptor for RSV, which is IGF1R^4^ and its coreceptor NCL^5^. RSV uses mitogenic signaling to initiate the entry process and regulate coreceptor trafficking. After entry into the cell, RSV is detected by pattern recognition receptors and we asked what other receptors might be used to propagate kinase signaling throughout virus replication, later in its replication cycle. In particular, Toll-like Receptor 7 (TLR7) detects the single-stranded RNA (ssRNA) produced as an RSV replication by-product^6,7^. TLR7 is expressed predominantly in antigen presenting cells like B cells, macrophages and dendritic cells and is present in late endosomes. *TLR7* transcripts have been detected in bronchial epithelial cells (BECs), the primary target of RSV infection^8^; however, the functional role of TLR7 in BECs has not been examined.

Here we show that TLR7 acts as a replication factor that supports optimal RSV replication in BECs through downstream activation of myeloid differentiation primary response 88 (MyD88) and extracellular signal-regulated kinase (ERK) MAP kinase pathway. Cellular localization studies identified TLR7 in the vicinity of RSV replication complexes in BECs. This led us to postulate that it was detecting RSV ssRNA produced by replicating RSV, resulting in ERK MAP kinase signaling that drives virus protein expression and suppresses cell death. We show a novel TLR7-ERK effector function that suppressed METTL3 ‘writer’ protein activity, reducing epigenetic RNA modifications. This METTL3 suppression resulted in further enhancement of TLR7 activation and RSV replication. Mice with a human TLR7 gain of function allele demonstrated enhanced RSV replication that caused an exacerbated disease phenotype in male mice, consistent with RSV disease prevalence and pathogenesis. Altogether our results suggest a novel role for TLR7 in RSV disease pathogenesis and the sex dependence of disease.

## TLR7 agonists increase RSV replication in bronchial epithelial cells

We tested a panel of commercially available nucleoside analogues for their effect on RSV infection in 1HAEo-BECs using a method we described previously^9^ (**Extended Data Fig. 1a**). This included the guanosine nucleoside analogues 7-allyl-7 8- dihydro- 8- oxo- guanosine triphosphate (loxoribine) and 1-b-D- Ribofuranosyl- 1,2,4- triazole- 3-carboxamide (ribavirin). Ribavirin was used as a control based on a well-defined *in vitro* and clinical profile, as an antiviral drug control^10^, and we observed inhibition of replication as expected^11^. However, there was a dose-dependent increase in RSV progeny production in the presence of TLR7 agonist, loxoribine, and CL097, a dual TLR7 and 8 agonist (**Extended Data Figs. 1b and c**), without any effect on the number of first round foci of infection (**Extended Data Fig. 1d**). Activating both TLR7 and TLR8 with CL097 made no significant difference to RSV replication compared to when TLR7 was activated alone by loxoribine. These results suggested that TLR7, alone, was responsible for enhancement of RSV replication caused by loxoribine in 1HAEo-BECs.

Recognition of viruses in the cytoplasm by TLR7 requires transport of cytosolic viral replication intermediates into the late endosome, and others have shown that the process of autophagy is required to mediate this process^12,13^. The process of autophagy occurs when the cell engulfs its cytoplasmic contents in membrane vesicles and it is required for optimal RSV replication^14,15^. To study autophagy during RSV infection in BECs we detected LC3 by immunofluorescence confocal microscopy. LC3 is a host protein that forms autophagic membrane vesicles and as it does it becomes detectable by confocal microscopy. We found that LC3 became visible during RSV infection and localized with RSV replication complexes as early as 3 hours post infection (**Extended Data Fig. 1e**). This suggested to us that RSV RNA could be accessible to TLR7 during infection.

**Extended Data Fig 1.**
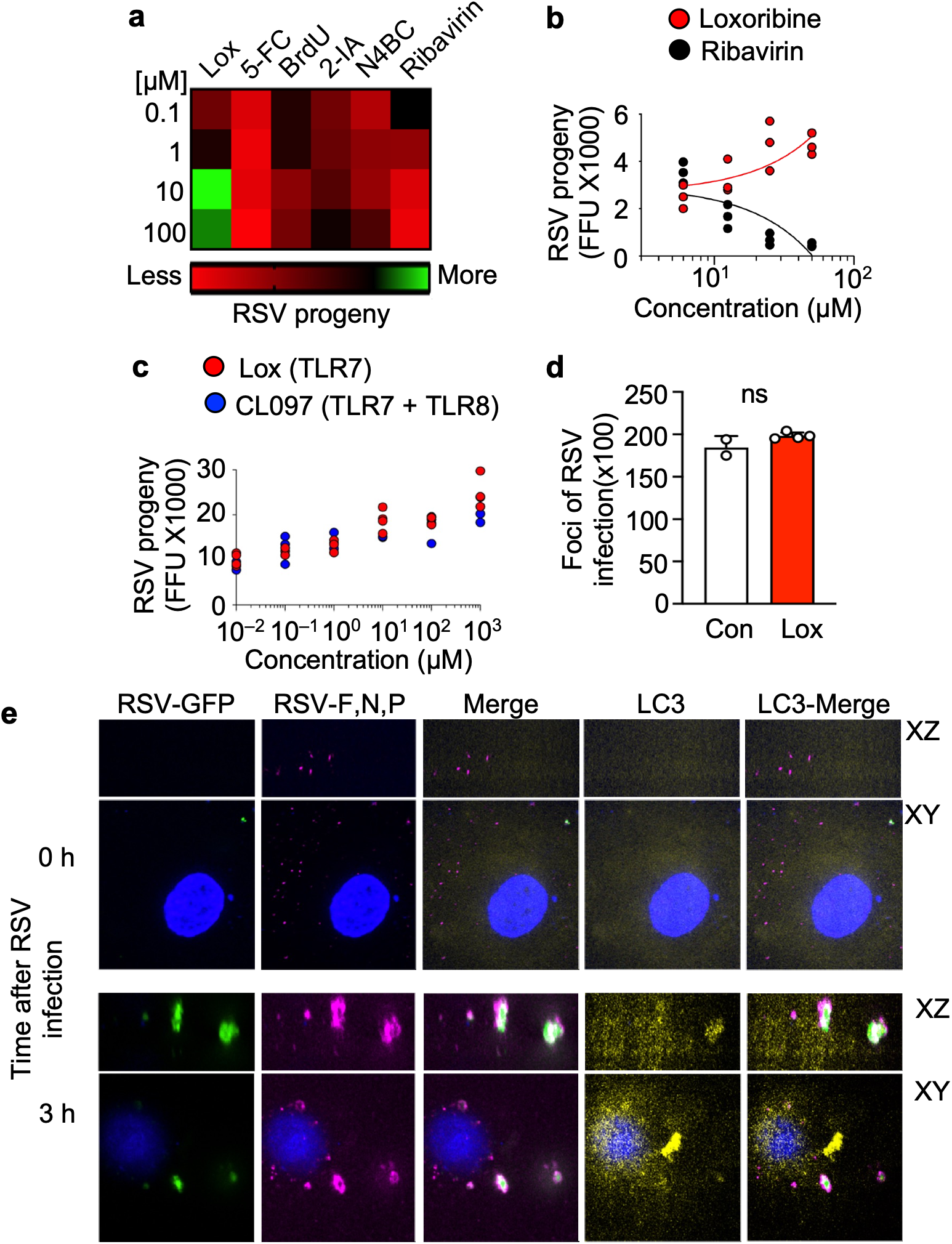
TLR7 agonists enhance RSV replication in human bronchial epithelial cells, **a,** RSV progeny release after nucleoside analogue treatment of lHAEo-cells with loxoribine (Lox), 5-fluorocytidine (5-FC), bromodeoxyuridine (BrdU), 2-iodoadenosine (2-IA), N4-benzyolcytidine (N4BC), and ribavirin from 0.1 - 100μM. **b-c,** RSV progeny release from lHAEo-cells treated with a concentration gradient of loxoribine, ribavirin or CL097, a TLR7 and 8 agonist, **d,** RSV infected lHAEo-cells quantified after treatment with Lox or DMSO control, **e,** Immunofluorescent images of RSV co­localization with LC3 at 0 and 3 hours post infection in primary BECs.

## *TLR7* is expressed in primary human bronchial epithelial cells

Given that loxoribine is a TLR7 agonist we explored whether TLR7 was expressed in BECs. The expression and function of TLR7 has been mainly studied in immune cells of myeloid origin like macrophages and dendritic cells, though it is also highly expressed in B cells^12,16^. TLR7 serves as an ssRNA pattern recognition receptor that detects the presence of ssRNA byproducts of viral replication. Macrophages and dendritic cells that detect ssRNA by TLR7 express high levels of type I interferons (IFN) and serve to present viral foreign antigens to T cells serving to initiate an immune response to a viral infection. However, the primary targets of RSV are ciliated BECs that are structural in nature and line the airways of the lungs. If TLR7 was playing a significant role during RSV infection we expected that TLR7 would be expressed in BECs.

To test whether TLR7 protein was expressed in primary human BECs we used monoclonal antibodies against the N and C terminal domains of TLR7 to detect it by western blot (**Extended Data Fig. 2a**) and with mass spectrometry analysis (LC-MS/MS) (**Extended Data Fig. 2b**). To broaden our analysis of TLR7 expression in the lung we employed an online internet resource of single cell RNA sequencing data derived from human lungs^17^. Here, we identified consistent *TLR7* expression in human lung epithelial cells, albeit not as plentiful as the *TLR7* expression observed in myeloid and lymphoid cells of the immune system^17^ (**Extended Data Figs. 2c** and **d**).

**Extended Data Fig 2.**
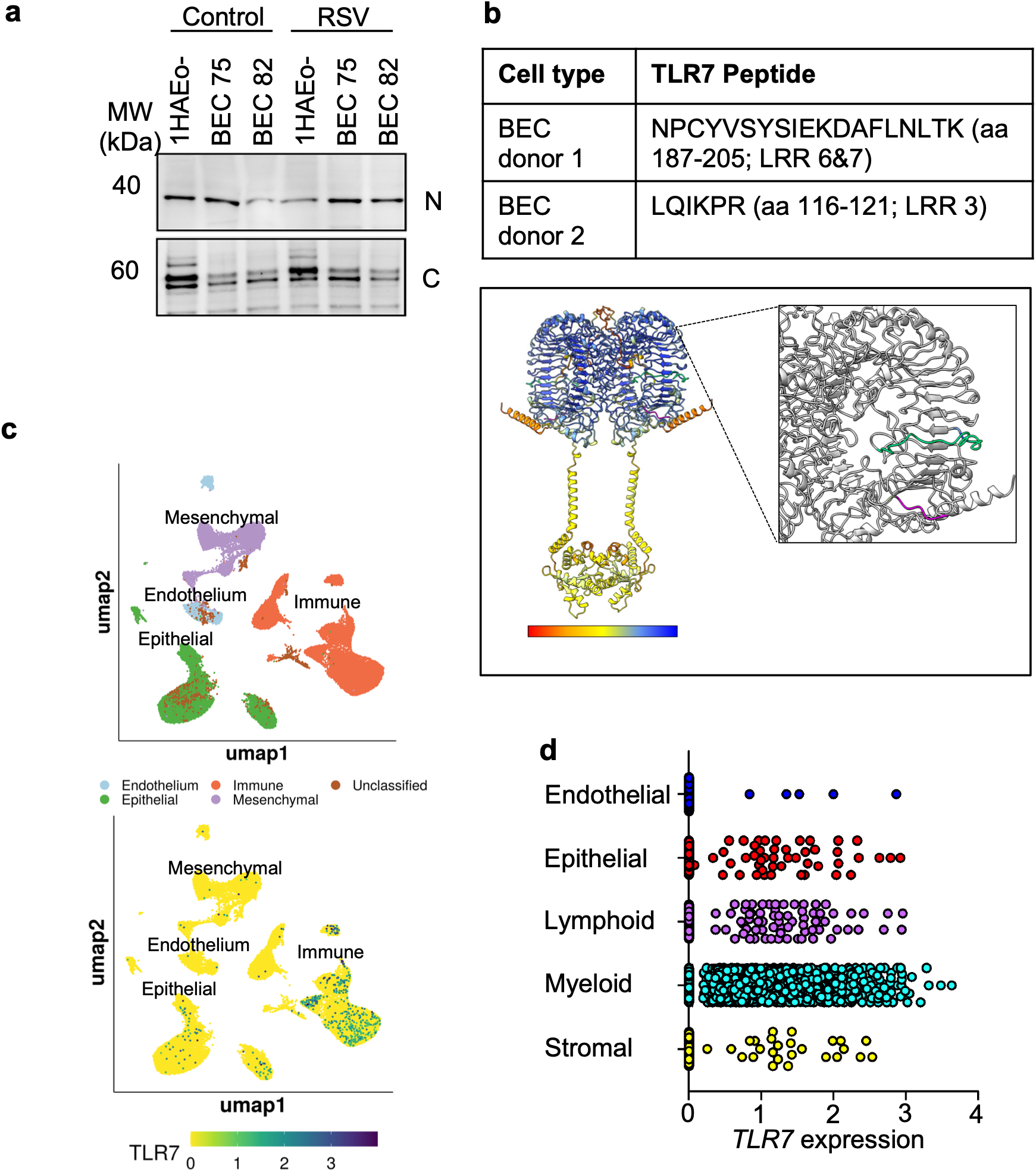
TLR7 is expressed in bronchial epithelial cells in human lungs, **a,** TLR7 N and C terminal fragments detected by western blot in uninfected and RSV infected immortalized lHAEo-cells and primary BECs from donors 75 and 82. Lysates were harvested at 24HPI. Blots are representative of 3 biological replicates, **b,** TLR7 peptides obtained from two human BEC donors by LC-MS/MS. Below is the predicted structure of TLR7 homodimer using Alphafold 3. **c,** Single cell sequencing shows human lung cellular compositions above, and TLR7 expression within these populations below. N=46500 cells, (Wang et al., eLife 2020). **d,** Relative TLR7 expression in different lung cell types determined by single cell RNA-seq. RNA was collected from donors aged 1 day to 33 years old with no underlying pulmonary disease. N=46500 cells, (Wang et al., eLife 2020).

## *TLR7* expression is sex-dependent in bronchial epithelial cells

The human *TLR7* gene is on the X-chromosome and therefore higher *TLR7* expression can be found in dendritic cells from females, which is due to higher X-chromosome gene dosage^18^. The *X-ist* RNA transcripts suppress the expression of genes from the silenced second copy of the X-chromosome in females by an epigenetic mechanism, however this suppression is less effective against genes further from the X-inactivation center locus like *TLR7*^19^.

We asked whether *TLR7* expression was higher in primary female BECs as compared to males. We found higher levels of *TLR7* mRNA expression in the two female BEC donors compared to the three male donors used in this study (**Extended Data Fig. 3a**), consistent with the higher *TLR7* expression observed in dendritic cells from female donors^18^. Additionally, we found that RSV infected female BEC donor cells produced more RSV progeny in untreated cells and in response to loxoribine treatment as compared to male BEC cells (**Extended Data Fig. 3b**).

Next, we surveyed the localization of TLR7 expression in 1HAEo- and primary BECs using confocal immunofluorescence microscopy (**Extended Data Figs. 3c** and **d**). Similar to what has been observed in myeloid and lymphoid cells^20^, we found that TLR7 in BECs was mostly confined to late endosome/lysosome compartments (**Extended Data Figs. 3d** and **e**). Pearson’s correlation analysis of the microscopy images showed that TLR7 colocalized most with vesicles in the cell that stained for LAMP1 late endosome marker and a Lysotracker dye that accumulates in lysosomes.

**Extended Data Fig 3.**
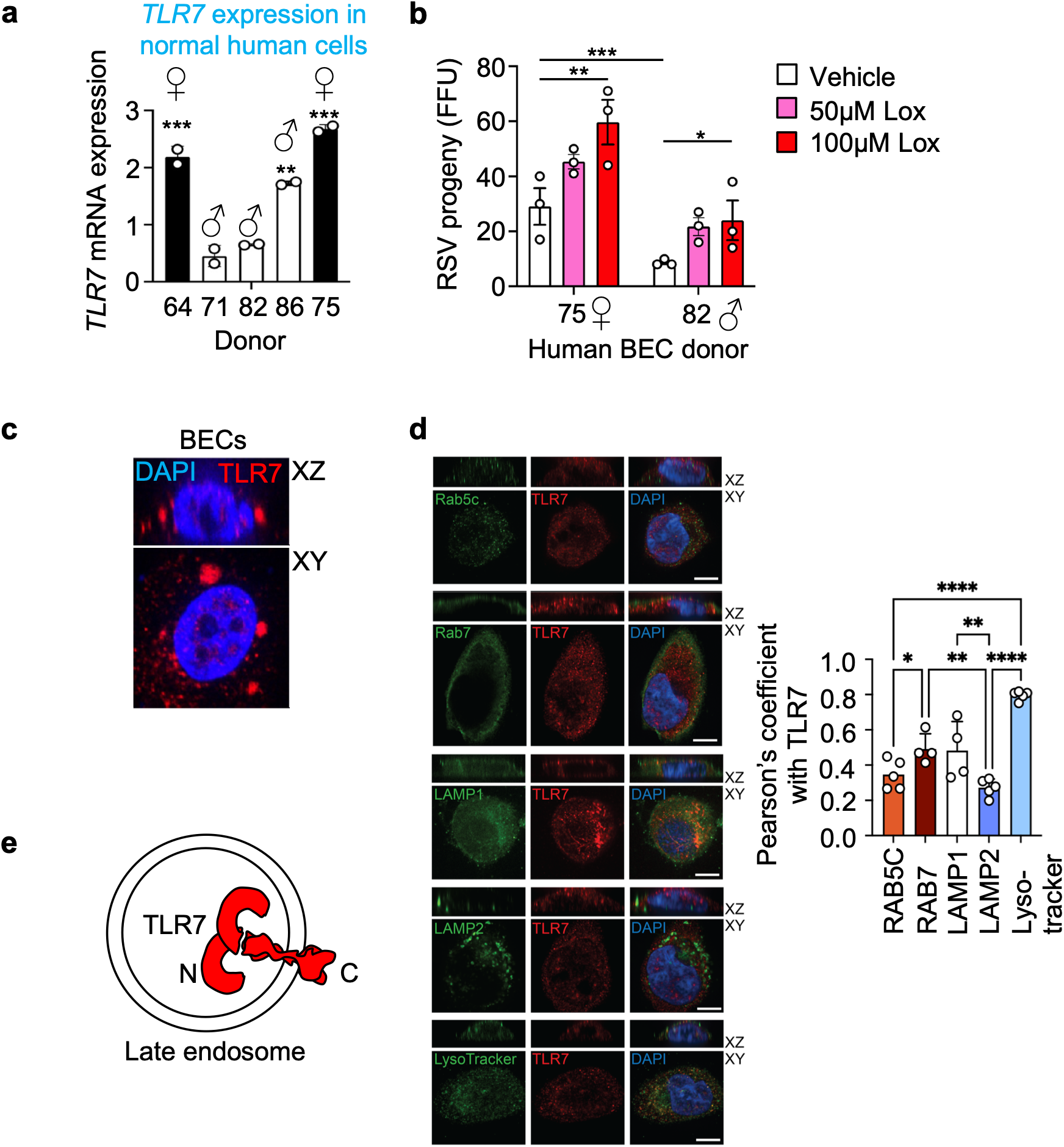
TLR7 localization in human bronchial epithelial cells. **a,** TLR7 expression in male and female human primary BECs determined by RT-qPCR relative to HPRT. N = 2 for each donor. **b,** RSV progeny release from female and male BEC donors treated with vehicle, 50μM, or 100μM ofloxoribine. Results are representative of3 biological replicates. **c,** Immunofluorescent TLR7 staining (red) in BEC primary cells also stained with DAPI (blue). Results are representative of 3 biological replicates. **d,** Pearson’s correlation coefficient ofTLR7 with endosomal markers determined from confocal immunofluorescent imaging in lHAEo- cells. N = 5 cells. Error bars= 8μm **e,** Schematic ofTLR7 positioning in a late endosome.

## TLR7 localizes to RSV inclusion bodies in BECs

A mechanism by which cells detect the presence of RNA from cytoplasmic RNA viruses like RSV is by autophagosome fusion with TLR7-containing late endosomes^12^. Due to LC3 expression during infection and its localization to RSV replication complexes (**Extended Data Fig. 1e**), and that TLR7 expression in BECs, we wanted to know whether TLR7 was also found near RSV replication sites. We found that GFP fluorescence from RSV-GFP occurs by 6-h after infection of primary BECs where GFP expression can be observed colocalizing with RSV F, N and P proteins in RSV inclusion bodies (**Extended Data Fig. 4a, Extended Data Fig. 1e**). In these experiments we stained for TLR7 as well and observed localisation to sites where RSV was producing GFP (**Extended Data Fig. 4a**) as well as the RSV-F glycoprotein (**Extended Data Fig. 4b** and **c**).

**Extended data Fig 4.**
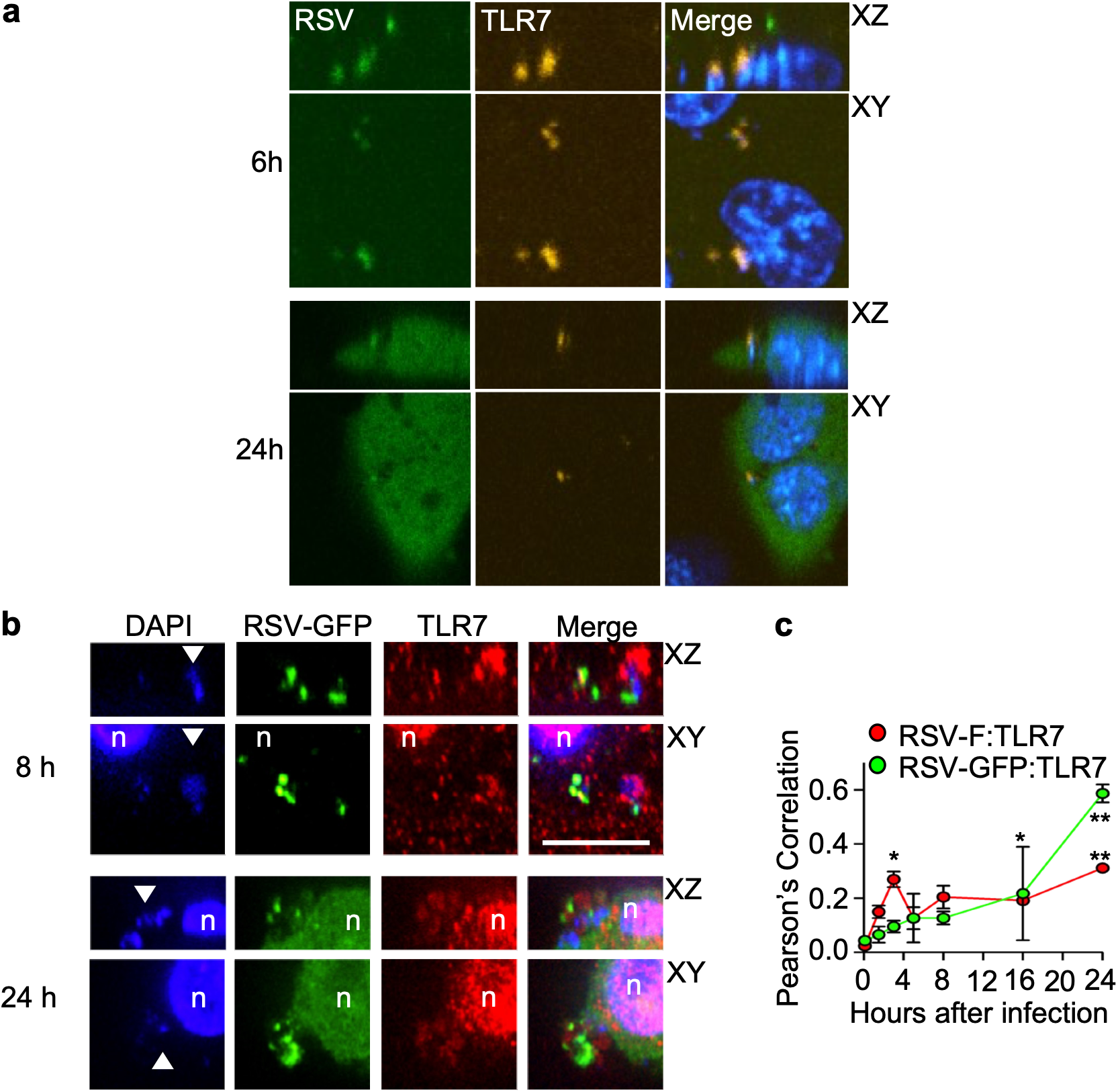
TLR7 associates with RSV replication complexes, **a,** RSV-GFP (green) and TLR7 (yellow) immunostaining at 6 and 24 hours following infection of primary BECs. **b,** RSV-GFP (green), DAPI (blue), and TLR7 immunostaining (red) in primary BECs. Nucleus (n) is marked, **c,** Pearson’s correlation coefficient between RSV-F and TLR7, and RSV-GFP at different points following RSV infection in determined from confocal imaging.

## TLR7 is involved in RSV replication after virus entry

We next studied the point that TLR7 was affecting in the RSV replication cycle. We postulated that TLR7 would act after RSV entry since TLR7 is typically an intracellular TLR found in late endosomes. In support of this, we found that there was no difference in the quantity of primary RSV foci in control or loxoribine treated 1HAEo-BECs (**Extended Data Fig. 1d**). This suggested to us that loxoribine was not facilitating virus entry into the cell because it did not increase the number of cells that became infected. We concluded from this that loxoribine was acting at a post-entry step.

To test when in the RSV replication cycle that TLR7 was acting, we added loxoribine at progressively later time points during RSV infection in 1HAEo-BECs and then measured the amount of progeny virus produced. The amount of RSV replication was greatest when loxoribine was added 8 hours after infection (**Fig. 1a**), that was about 5 hours after the onset of RSV protein expression (**Extended Data Fig. 1e**). Altogether these data suggested to us that TLR7 was acting after RSV had entered the cell and started to replicate.

**Figure 1.**
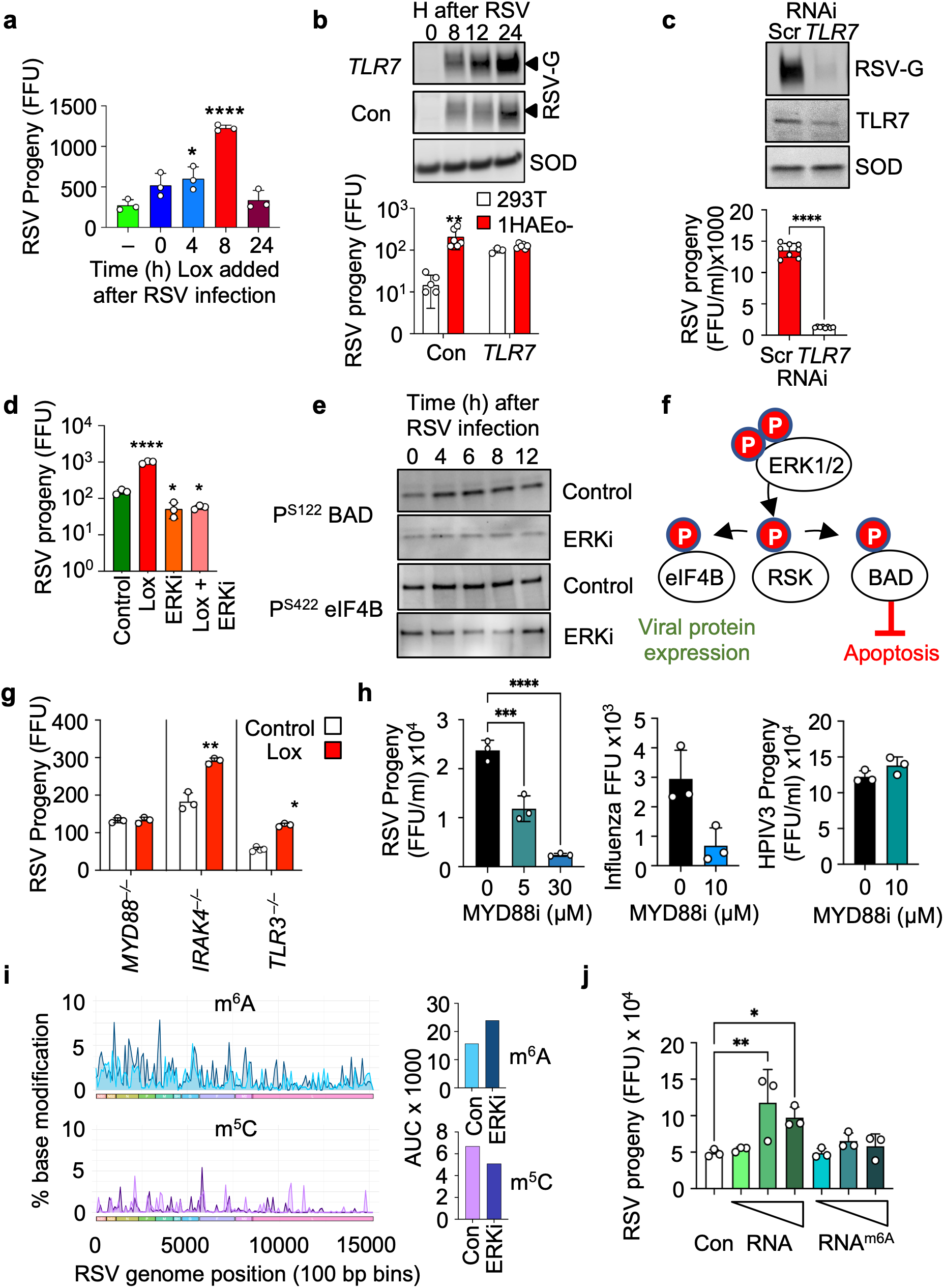
TLR7 signaling is required for optimal RSV replication, **a,** RSV progeny release in focus forming units (FFU) from lHAEo-cells treated with 200pM loxoribine (Lox) at 0, 4, 8, or 24 hours post infection (hpi). **b,** RSV progeny release from HEK293T and lHAEo-cells transfected with either control plasmid or TLR7 expression plasmid. Above, a western blot shows RSV-G protein levels at different times following RSV infection in HEK 293T cells transfected with control plasmid or TLR7 expression plasmid. SOD1 protein expression is shown as a loading control, **c,** RSV progeny release from lHAEo-cells treated with scramble (Scr) or TLR7 siRNA. Above is a western blot showing RSV-G and TLR7 protein expression in lHAEo-cells treated with Scr or TLR7 siRNA, with SOD1 protein expression as a loading control, **d,** RSV progeny release from lHAEo-cells treated with 200pM Lox, 10pM of U0126 (ERKi), or both at 4hpi. **e,** Western blot of phosphorylated BAD and elF4B in lHAEo-cells that were untreated (control) or treated with lOpM of U0126 (ERKi) and harvested at 0, 4, 6, 8, and 12 hours post RSV infection, **f,** A model of signalling mediator phosphorylation following RSV infection, **g,** RSV progeny release from *MyD88, IRAK4,* and *TLR3* null fibroblast cells treated with 200pM Lox or DMSO at 4hpi. **h,** Progeny RSV and HP1V3 release from lHAEo-cells treated with MyD88 inhibitor TJ-M2010-5 (MyD88i) at 4hpi is shown on the right and left. In the centre, influenza A infected A549 cells in FFU were treated with MyD88i at 2hpi and quantified at 48hpi. **i,** Percentage of N^6^-methyladenosine (m^6^A) and 5-methylcytosine (m^5^C) base modifications averaged over 100bp bins over the RSV genome in RNA harvested from infected primary BECs treated with DMSO or U0126 (ERKi). Position of RSV genes from NS1 to L are depicted below the graphs. Area under the curve (AUC) from each treatment group quantifies the results from these graphs. Results are from one primary donor, **j,** RSV progeny release from lHAEo-cells treated at 4hpi with either TE control or 100nM, 500nM, and 1μM unmodified RNA or m6A modified RNA. **a,b, d-h,** and **j** are representative of 3 biological replicates, **c** is representative of 2 biological replicates.

## TLR7 is required for optimal RSV replication in bronchial epithelial cells

To determine whether TLR7 was required for optimal RSV replication we sought to modify the expression of TLR7 in the absence of TLR agonists. We expressed *TLR7* in 1HAEo-BECs and in 293T embryonic kidney cells, a cell line that does not normally express *TLR7* (**Fig. 1b)**. Here we found that RSV replication was enhanced in 293T cells to the same level as we saw in the 1HAEo-BECs when *TLR7* was overexpressed. Conversely, silencing *TLR7* gene expression in 1HAEo-BECs significantly reduced RSV replication (**Fig. 1c**). In summary, RSV replication required *TLR7* expression in human BECs for optimal replication.

## Optimal RSV and influenza replication are ERK MAP kinase dependent

We and others have shown that RSV replication is enhanced by host cell MAP kinase activation^21,22^ acting to regulate the expression of RSV genes^21^ and glycoprotein trafficking^23^. We confirmed that in 1HAEo-BECs, RSV progeny release can be reduced by a MEK inhibitor (UO126) and an ERK inhibitor (SCH772984) (**Extended Data Fig. 5a**).

Most TLRs, including TLR7, can activate host cell MAP kinases when stimulated^24,25^. From these prior studies we postulated that RSV uses TLR7 activation to trigger the ERK MAPK pathway to drive optimal RSV replication (**Extended data Fig.5b**). ERK MAPK was rapidly activated following RSV infection and there was consistently higher phosphorylation during a 24 hour time course of RSV infection (**Extended Data Fig. 5c**). Conversely, ERK MAPK inhibition blocked loxoribine-dependent enhancement of RSV replication in 1HAEo-cells (**Fig. 1d**). Combined treatments with loxoribine and ERK MAPK inhibitor (MEK1/2 inhibitor UO126) were non-toxic to the BECs (data not shown). These results suggested that loxoribine enhances RSV replication by triggering ERK MAPK signaling activity.

**Extended Data Fig 5.**
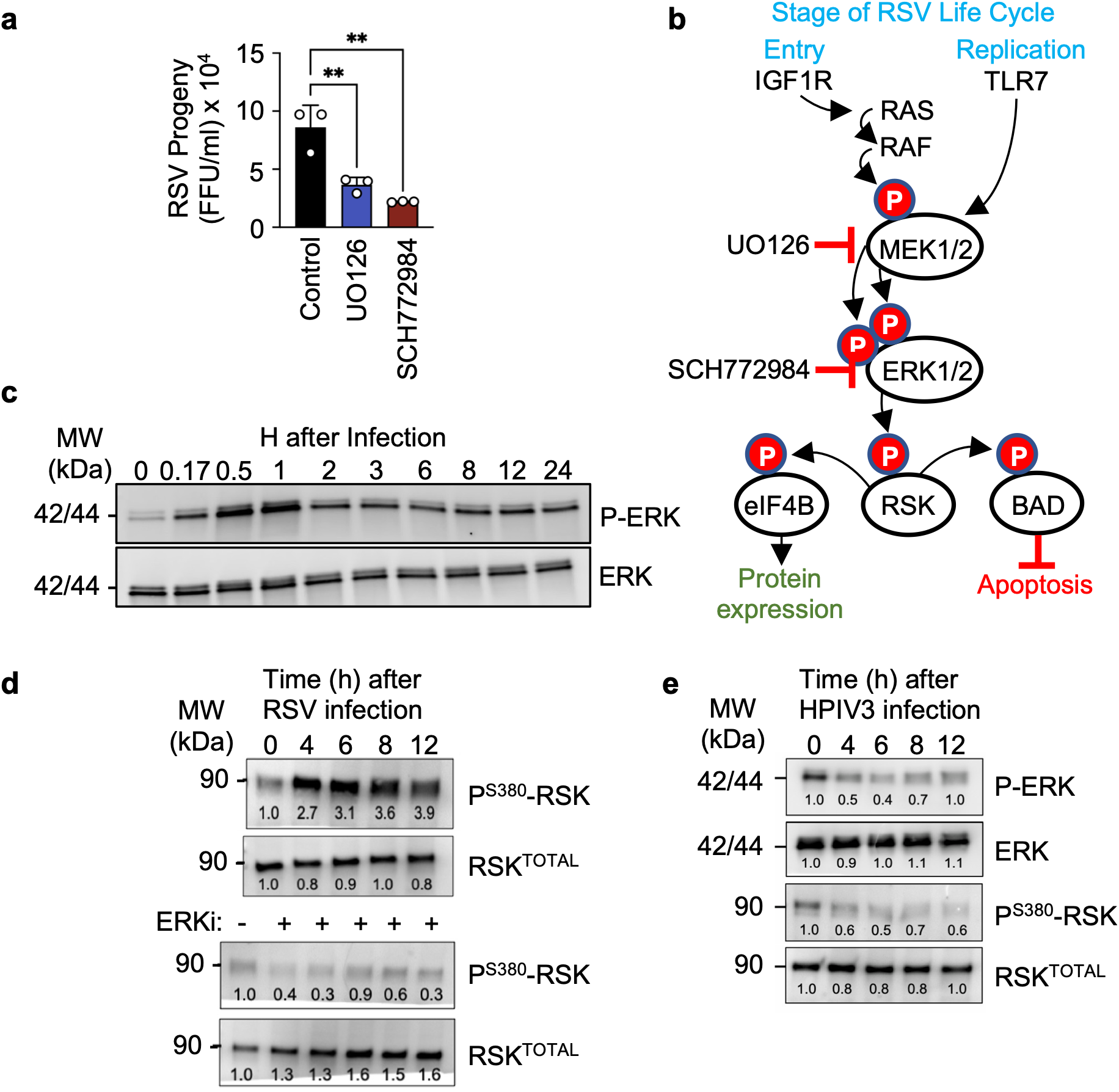
MAPK pathway signalling is activated by RSV infection, **a,** RSV progeny release from lHAEo-cells following U0126 and SCH772984 treatment at 4hpi. **b,** A schematic of common signalling at different points in the RSV life cycle, **c,** ERK phosphorylation following RSV infection in lHAEo-cells is shown by western blot, **d,** p90RSK phosphorylation at S380 in lHAEo-cells that were untreated (top) or treated with U0126 (ERKi, bottom) and infected with RSV is shown by western blot. Densitometry relative to loading controls is shown below each band, **e,** ERK and p90RSK phosphorylation in lHAEo-cells infected with HPIV3 are shown by western blot. Densitometry relative to loading controls is shown below each band **a, c,** and **d** are representative of 3 independent experiments.

## RSV infection activates p90 RSK in BECs

The dependence of RSV replication on ERK MAP kinase activation has been established^21–23^, but the downstream effectors of ERK during RSV infection have not been studied in detail. ERK MAP kinase acts to support cell survival by regulating functions such as protein expression, proliferation, and apoptosis^26,27^. We asked whether ERK played these roles during RSV infection because RSV infected cells are seemingly apoptosis resistant (reviewed in:^3^), and viral protein expression is dependent on TLR7 signaling (**Figs. 1b** and **c**). We tested a downstream effector of ERK called p90 RSK which has roles in regulating protein expression and cell survival. We noted that phosphorylation of S380 on p90 RSK was observable for the duration of RSV infection like ERK MAPK (**Extended Data Fig. 5d**). S380 phosphorylation was also blocked with a MEK inhibitor. Similar to what we have found for RSV, it has previously been shown that late activation of ERK MAPK and its downstream targets like p90 RSK by influenza A infection is essential for optimal replication^28^. To determine whether this signalling pathway activation was common to other respiratory viruses, we tested whether HPIV3 infection caused similar phosphorylation (**Extended Data Fig. 5e**). Contrarily, we found that HPIV3 infection led to a decrease in ERK and RSK phosphorylation.

Effectors directly phosphorylated by p90RSK, such as BAD, which has an anti-apoptotic effect, and eIF4B, which has a pro-translational effect, were activated upon RSV infection and had a decrease in phosphorylation following MEK inhibitor treatment (**Fig. 1e** and **f, Extended Data Fig. 5b**).

## Optimal RSV replication is MyD88 dependent

All TLRs, except TLR3, signal through a TLR-specific intracellular signaling adaptor called MyD88 and interleukin-1 receptor-associated kinase 4 (IRAK4)^25^. We found that both *MYD88* and *IRAK4* are normally expressed in two donors of human primary BECs (**Extended Data Fig. 6**) where *MYD88* expression was significantly elevated (**Extended Data Fig. 6b**) but IRAK4 was significantly reduced (**Extended Data Fig. 6c**) after 48 hours of RSV infection. These data may suggest that RSV has evolved to enhance *MYD88* expression to support RSV replication while evolving to suppress *IRAK4* expression to counter the antiviral effects of IRAK4. *IRAK4* expression was similar to uninfected cells when TLR7 activity was inhibited, suggesting that suppression of *IRAK4* expression is TLR7 dependent during RSV infection of BECs. In contrast, *MYD88* expression remained significantly elevated even after TLR7 inhibition (**Extended Data Fig. 6b**). Additionally, we found that inhibition of TLR7 activity led to reduced overall differential gene expression upon RSV infection, particularly in interferon stimulated and pattern recognition receptor associated genes (**Extended Data Figs. 6a-e**).

We postulated that if TLR7 signaling was associated with RSV replication, the involvement of MyD88 would be expected and, consequently, there should be no response to loxoribine in *MYD88^−/−^* cells. We used *TLR3^−/−^* cells as a control because TLR3 has been shown not to have an effect on RSV replication^29^. Using human *MYD88^−/−^, IRAK4^−/−^*, and *TLR3^−/−^* fibroblast cells^30,31^, we previously showed that MyD88, IRAK4 and TLR3 were not required for RSV entry^4^. To study if virus replication was affected in these cells, we quantified progeny virus produced during RSV replication from the *MYD88^−/−^*, *IRAK4^−/−^* and *TLR3^−/−^* cells. We noted that there was a significant increase in RSV progeny release during TLR7 agonist (loxoribine) treatment of *IRAK4^−/−^* and *TLR3^−/−^* cells (**Fig. 1g**). Others have shown previously that IRAK1 and 2 can propagate MyD88 signaling in the absence of IRAK4^32^. Importantly we noted that RSV replication was not influenced by TLR7 agonist treatment in *MYD88^−/−^*cells.

To further test the involvement of MyD88 in our model, we asked whether MyD88 signaling inhibitors could suppress RSV replication. We also tested influenza because it uses ERK MAP kinase signaling to export vRNPs from the nucleus^33^, so we hypothesized that it may also use MyD88 signaling to trigger this pathway. We found that HPIV3 did not activate ERK MAP kinase so we used it as a negative control (**Extended Data Fig. 5e**). We observed reduction of RSV progeny release and influenza infection of human A549 alveolar cells by treatment with MyD88 inhibitors (**Fig. 1h**). HPIV3 infection was not affected by MyD88 inhibition. Together, these results suggested to us that MyD88, and not IRAK4 or TLR3, are required for optimal RSV and influenza replication and infection through TLR7 stimulation.

**Extended Data Fig 6.**
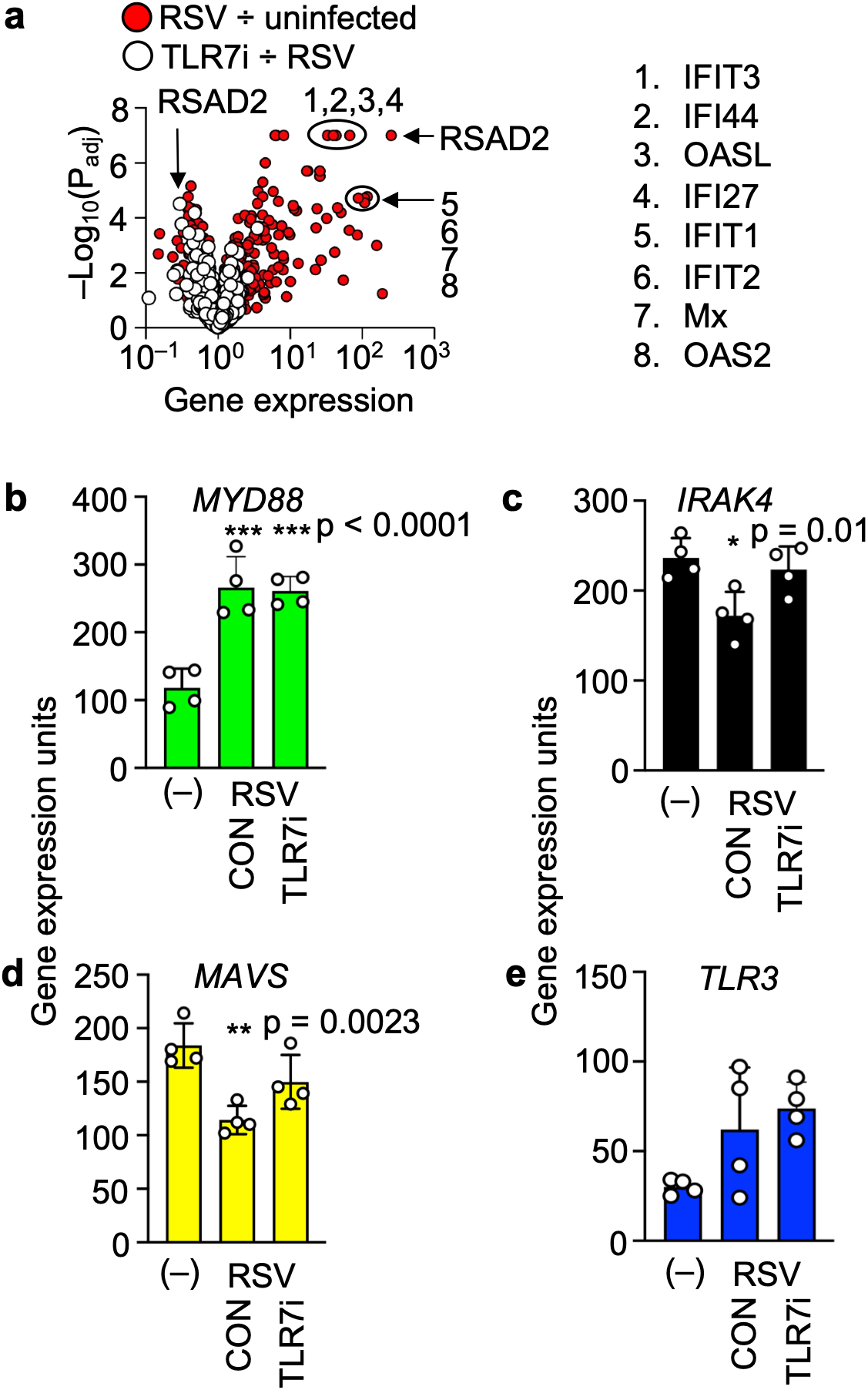
TLR7 inhibition affects the immune response to RSV infection, **a,** Gene expression in primary BECs treated with IRS661 (TLR7i) or untreated (RSV) following 48 hours of RSV infection. In red, gene expression in RSV infected cells is compared to uninfected cells. In white, gene expression in TLR7 inhibitor infected cells are compared to untreated infected cells. Gene expression was determined using Nanostring nCounter Analysis. N = 2 x 2 donors per replicate, **b-e,** Gene expression from specific immune genes of interest in uninfected, untreated control, and IRS661 treated (TLR7i).

## RSV ERK signaling reduces m^6^A modification of RSV mRNA

Epigenetic modification of viral RNA by host enzymes is emerging as an important aspect of viral pathogenesis. N^6^-methyladenosine (m^6^A) modifications are one of the most prevalent and abundant RNA modifications in the cell^34,35^. We read with interest that ERK MAPK signaling can affect RNA base modification by stabilizing the METTL3-METTL14 writer complex that adds methyl groups to adenosine at position 6^36^. These modifications can impact many facets of the viral replication cycle, including transcript stability, translation, and immune evasion. There are conflicting reports however about how N^6^-methyladenosine (m^6^A) modifications affect RSV infection, possibly due to differences in the cell types used in the studies^37–40^. This could be important in the design and administration of mRNA vaccines which are gaining rapidly in popularity^41^.

To address the roles of the METTL epigenetic writer enzymes in RSV replication, we first asked whether ERK MAPK inhibition could affect the base modifications of RSV in BECs. To do this, we sequenced RSV mRNA from RSV infected 1HAEo-cells and detected 5-methylcytosine (m^5^C) and m^6^A modifications in the presence and absence of an ERK inhibitor (**Fig. 1i**). Here we found an overall increase in m^6^A of RSV genome transcripts but a decrease in m^5^C in ERKi treated BECs. From this data we posited that RSV may have evolved to decrease m^6^A by ERK activation in BECs to reduce the ‘self’ appearance of its RNA to TLR7^42^, thereby increasing TLR7 RNA detection and signaling. By our working model, the reduction of m^6^A modification by ERK activation during RSV replication would enhance TLR7 recognition and signaling.

To test this, we next asked whether treatment of BECs with unmodified and m^6^A modified RNA oligonucleotides could activate TLR7, and increase RSV replication, like a TLR7 agonist. We treated 1HAEo-BECs with m^6^A modified and unmodified RNA oligonucleotides 4 h after RSV infection. Viral progeny produced by the cells were detected as an indicator of viral replication (**Fig. 1j**). We found that control RNA oligonucleotides significantly increased RSV replication, whereas there was no increase in RSV replication by the m^6^A modified RNA oligonucleotides.

## Human samples from females have higher RSV viral loads than males

We postulated that females would have higher viral loads than males given our in vitro results (**Extended Data Fig. 3a** and **b**). We have previously reported on the phylogenetics of RSV in western Canada compared to RSV worldwide^43^. Samples in this study were obtained from patients by nasopharyngeal (NP) swabs from which RSV viral loads were measured by qRT-PCR for comparison using a standard curve to determine equivalent focus forming units^43^. Reanalysis of this data set showed that females had significantly higher viral loads than males over all age groups (**Extended Data Fig. 7a**) and when stratified into those one year of age and younger (**Fig 2a**). This stratified age group had the highest statistical power and is the one most likely to seek medical attention due to RSV infection. To study *TLR7* expression in these samples, we determined *TLR7* mRNA levels, and we found that their RSV viral loads were correlated with *TLR7* expression in both sexes (**Fig. 2b**). We marked the range of the x-axis of the graph for males on the graph of female swabs to highlight the difference in *TLR7* expression.

**Figure 2.**
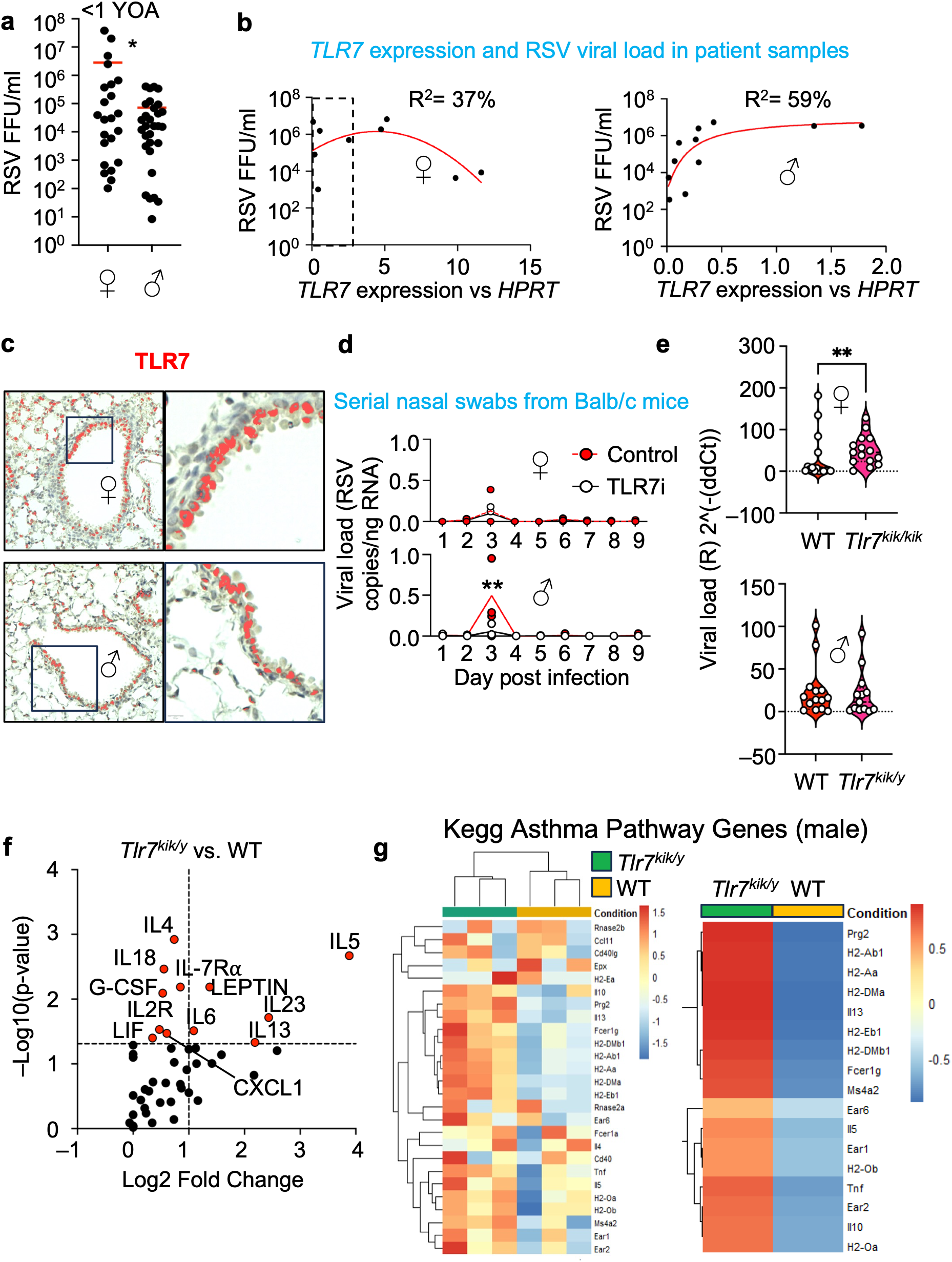
TLR7 is an RSV pathogenesis factor in vivo, **a,** Sex differences in RSV viral load in nasopharyngeal (NP) swabs from infant <1 year of age (YOA). RSV titres were determined using RT-qPCR results and focus forming units (FFU) per mL were calculated by comparing RSV cycle numbers to a standard curve, **b,** RSV viral load from male and female NP swabs compared to TLR7 expression levels. Using RT-qPCR, RSV RNA was quantified as described in **a,** and TLR7 RNA expression was determined relative to housekeeping gene hypoxanthine-guanine phosphoribosyltransferase (HPRT). Non-linear regression was performed using Graphpad prism software. Female swabs (right) and male swabs (left) are shown seperately. The dotted box on the female graph represents the range of the x-axis of the male graph, **c,** TLR7 histology staining in Balb/c mice lungs. TLR7 in lung sections was immunostained using HRP detection and hematoxylin counterstained. QuPath Software was used to mark TLR7 staining in red. Images are representative of 4 biological replicates for males and females, **d,** RSV viral loads in RSV infected Balb/c female (above) and male (below) mice who were treated with IRS661 (TLR7i) or control oligodeoxynucleotide (ODN). Nasal-oral swabs were collected each day, and viral loads were determined by RT-qPCR in comparison with a standard curve and normalized to ng of input RNA. N = 3 mice for each treatment in both males and females, **e,** RSV viral load in female homozygous kika mice *(Tlr7^kik/kik^)* (above) and male hemizygous kika mice *(Tlr7^kik/y^)* (below) as compared to WT mice of the corresponding sex. Lung RNA was harvested 3 days post RSV infection and viral load was determined using RT-qPCR as R value (=2^-(dCt-10)^) relative to HPRT. Mice that did not have a detectable viral load were excluded, **f,** Volcano plot of plasma cytokine protein measured as net mean fluorescent intensity (MFI) from RSV infected *Tlr7^kik/y^* mice compared to WT males. 48 cytokines were assayed. P-value was determined using wilcoxon rank-sum test. N = 13 for each genotype. Red points have a p-value >0.05 **g,** Lung RNA-seq results for KEGG asthma gene set (mmu05310) in RSV infected *Tlr7^kik/y^* mice compared to WT males. Heatmap colours represent Z-score values. N = 3 for each genotype. The left heatmap shows individual replicates, while the right heatmap shows genotype averages for a subset of genes.

**Extended data Fig 7.**
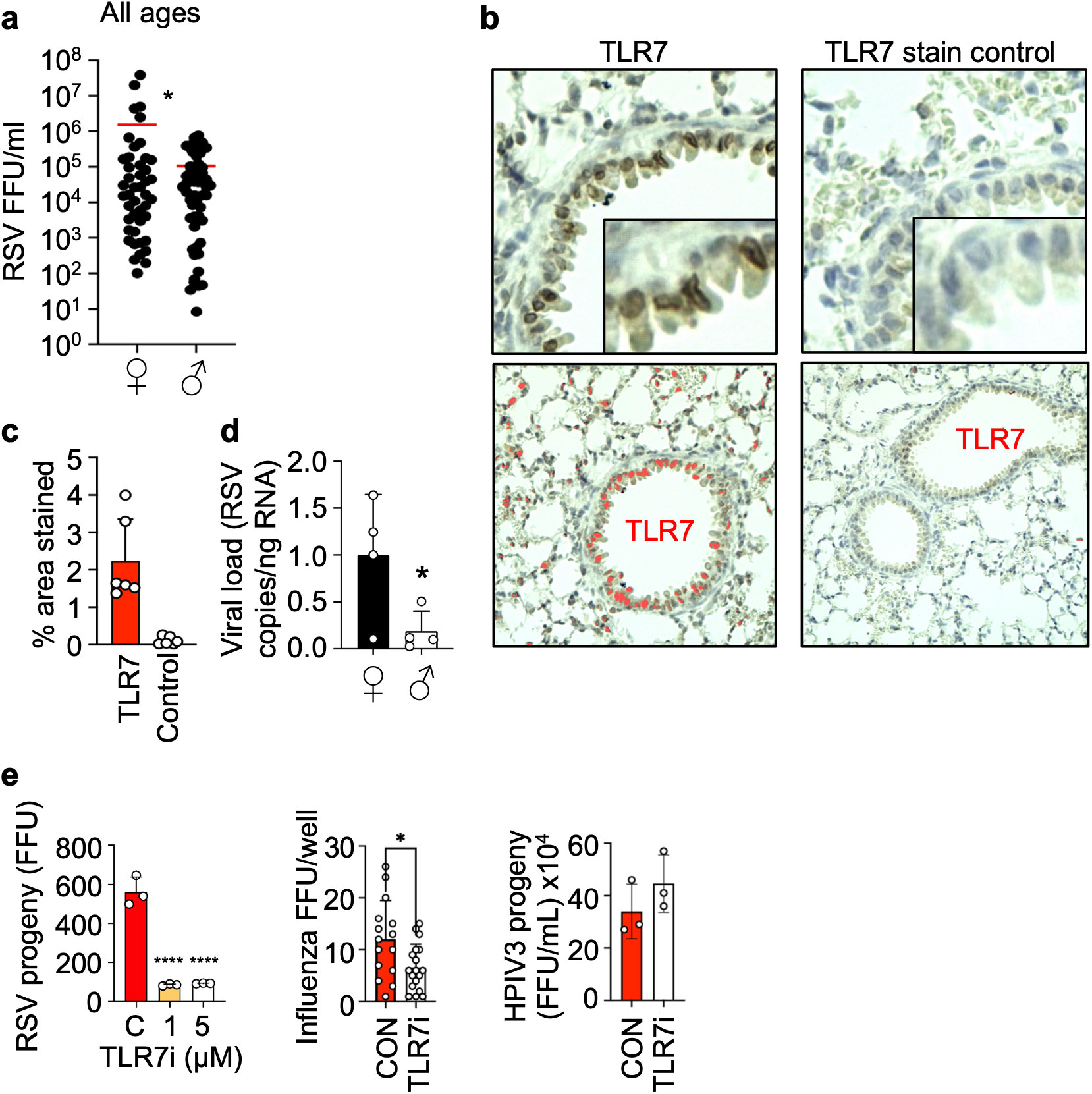
Sex differences in RSV infection, **a,** RSV viral loads from nasopharyngeal swabs RSV titres were determined using RT-qPCR results and focus forming units (FFU) per mL were calculated by comparing RSV cycle numbers to a standard curve, **b,** TLR7 immunohistology staining in tissue from a balb/c mouse (TLR7) compared to tissue with recombinant TLR7 protein added (TLR7 stain control). TLR7 staining is marked in red by QuPath software, **c,** Quantification of **b.** using QuPath software, **d,** RSV viral loads in female and male Balb/c mice 5 days post infection. Viral load in mouse lung RNA was determined by RT-qPCR in comparison with a standard curve and normalized to ng of input RNA. N = 4 for both males and females, **e,** RSV progeny release from lHAEo-cells treated with control ODN (C), 1, or 5pM of IRS661 (TLR7i) (left). Influenza infected A549 cells quantified at 48hpi after treatment with IRS661 or control ODN (con) at lpM at 2hpi (centre). HPIV3 progeny release from lHAEo-cells treated with IRS661 or control ODN (con) at lpM at 2hpi (right).

## TLR7 is expressed in the airways of mice

There have been previous studies of RSV infection in the Tlr7^−/−^ mouse model^7^. Here, they reported increased inflammation in the Tlr7^−/−^ mice compared to the WT matched controls due to lack of control of infection. Consistent with our results, they also reported higher viral loads in female mice compared to males and reduced RSV viral loads in Tlr7^−/−^ mice compared to their WT counterparts, supporting the role of TLR7 during RSV replication that we report here.

To determine whether TLR7 was expressed in the airways of Balb/c mice, we detected TLR7 by immunohistochemistry of Balb/c mouse lungs (**Fig. 2c**, **Extended Data Figs. 7b** and **c**). To rule out unspecific detection of TLR7 we neutralized the staining by the primary antibody against TLR7 using recombinant TLR7 protein, which we added to the tissue during the staining process. Stain detection software detected TLR7 in the BECs lining the airways, as well as in type II pneumocytes and alveolar macrophages, but not in the TLR7 staining control (**Extended Data Fig. 7c**). Consistent with what we found in human samples (**Extended Data Fig. 7a**), we found that viral loads in male Balb/c mice are significantly lower than in female mice (**Extended Data Fig. 7d**).

Prior studies of TLR7 and RSV in mice sampled infected tissue 5-7 days after inoculation with virus^7^. We surveyed when RSV was being shed from the nares of RSV infected Balb/c mice in the presence of a TLR7 inhibitor and controls (**Fig. 2d**). Here, we detected peak nasal shedding of RSV 3 days after infection. There was also significant inhibition of RSV in the lungs of male mice treated with TLR7 inhibitor, but not in the female mice. The effect of TLR7 inhibition in male mice during RSV infection resembles what we see in cells. In 1HAEo-BECs, TLR7 inhibitor decreases RSV progeny release as well as influenza infection in A549 cells but does not affect HPIV3 progeny release (**Extended Data Fig. 7e**). We believe that the effect of TLR7 inhibition on RSV infection in males only reflects the higher expression of *Tlr7* in females. This suggests that sufficiently high TLR7 expression helps to drive RSV replication but may simultaneously have downstream protective roles against RSV pathogenesis.

## Gain-of-function *TLR7* mice have more RSV infection

There are rare *TLR7* gain-of-function variants in the human population that cause auto-immune diseases like systemic lupus erythematosus^44,45^. To test the role of TLR7 as an RSV replication factor we infected mice engineered to express the gain-of-function *TLR7*^Y264H^ gene (*kika*) variant. These mice begin to develop lupus-associated antinuclear antibodies at 6 weeks of age^44^, and overall mortality is highest in female homozygous mice (*Tlr7*^kik/kik^), followed by male *Tlr7*^kik/y^ mice, and is lowest in female heterozygous mice (*Tlr7*^kik/+^) (**Extended Data Fig. 8a**). We infected *kika* and littermate control mice at 4-5 weeks old, before onset of disease. From the work we present here thus far we postulated that this gain-of-function *kika* mouse model would have higher levels of RSV replication compared to their WT counterparts. We infected the mice with RSV and sacrificed them 3 days later consistent with the peak of viral shedding (**Fig. 2d)**. Viral loads were significantly higher in the female mice homozygous for the *kika* allele (**Fig. 2e**), but there was no difference in viral loads between the male *Tlr7*^kik/y^ and male WT mice.

**Extended data Fig 8.**
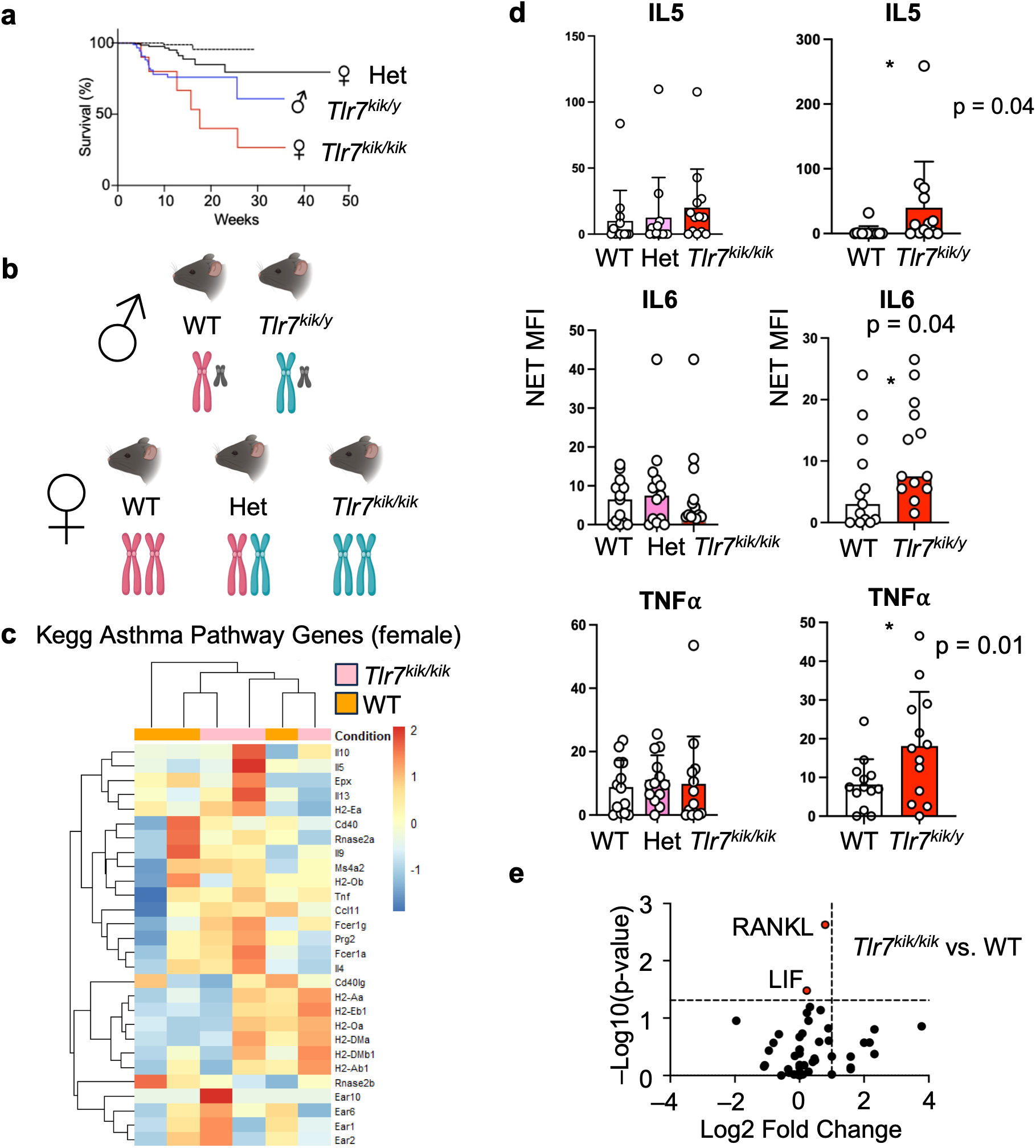
Cytokine and gene expression responses to RSV infection in TLR7 gain-of-function mice, **a,** Kaplan-Meier curve of *kika* mouse survival by genotype, **b,** Schematic of female and male *kika* genotypes, **c,** Lung RNA-seq results for KEGG asthma genes (mmu05310) in RSV infected *Tlr7^kMklk^* mice compared to WT females. Heatmap colours represent Z-score values. N = 3 for each genotype, **d,** IL-5, IL-6, and TNFoc protein expression in RSV infected female (left) and male (right) mouse plasma. Protein is quantified by net mean fluorescent intensity (MFI) and mouse genotypes are shown on the x-axis. N = 13 for each genotype, **e,** Volcano plot of plasma cytokine net MFI from RSV infected *Tlr7^k,k/kik^ mice* compared to WT females. 48 cytokines were assayed. P-value was determined using wilcoxon rank-sum test. N = 13 for each genotype. Red points have a p-value >0.05

## Male mice with a gain-of-function *TLR7* express higher levels of RSV disease markers

We next asked what effect the *kika* allele would have on expression of RSV disease markers in homozygous female *Tlr7*^kik/kik^ and male *Tlr7*^kik/y^ mice (**Extended Data Fig. 8b**). The patterns of expression of cytokines and chemokines in the lungs are an indicator and cause of severe disease outcomes, leading to narrowed and clogged airways^46^. In particular, IL-5-mediated attraction of eosinophils into the lungs causes hypersensitivity in the airways^47^. This is a target of a licensed monoclonal antibody drug that is used to treat eosinophilic asthma^48^. To understand the earlier responses to RSV replication we detected cytokine and chemokines protein expression in plasma 3 days post infection. A cytokine array showed that IL-5 and a skewing to a type 2 inflammatory response were significantly elevated in male *Tlr7*^kik/y^ mice, markers of severe RSV disease and asthma in mice and humans^49^ (**Fig. 2f, Extended Data Fig. 8d**). This was accompanied by significantly higher TNFα in the male *Tlr7*^kik/y^ mice, compared to their WT counterparts (**Extended Data Fig. 8d**). However, in female *Tlr7*^kik/kik^ mice, we observed minimal change in cytokine production when compared to WT female mice (**Extended Data Fig. 8e**). Due to the association of IL-5 and RSV infection with asthma, we examined the expression of asthma genes in both male genotypes by bulk RNA-seq of infected lungs. We found that male mice with the *Tlr7*^kik/y^ allele had higher expression of genes involved in asthma (**Fig. 2g**). This pattern was not consistent in the female mice with the *Tlr7*^kik/kik^ allele (**Extended Data Fig. 8c**).

## Discussion

Here we show that TLR7 is a core factor that drives RSV replication and disease pathogenesis, and we believe that this is one of the most highly detailed studies of TLR7 outside of immune cells. We show that TLR7 is normally expressed in BECs and detects RSV replication complexes in BECs. The resulting ERK MAP kinase activation drives optimal RSV replication. To determine whether the effect was common in other respiratory virus infections we tested HPIV3 and influenza A virus. TLR7 and MyD88 inhibition had no effect on HPIV3 replication and progeny release, however, we found that influenza infection was promoted by TLR7 signaling in airway epithelial cells. Others have found that TLR7 and RIG-I in responding monocyte derived dendritic cells are required for optimal influenza infection^50^. RSV and influenza infections provide an example of how inflammation and viral replication are part of a positive feedback loop, an emerging paradigm in viral infection^51^. The downstream effectors of ERK MAP kinase enhance viral protein synthesis and suppress cell death mechanisms that would otherwise halt virus infection. In male mice we found that eosinophilic asthma disease markers were significantly elevated in male mice with a gain of function *TLR7* gene, compared to the WT controls. This is consistent with human markers of severe disease, and higher risk of asthma development following RSV infection, a phenomenon that is more prevalent in males^52,53^. Preventing TLR7 expression or inhibiting its activation significantly reduced RSV replication and spread in cells and male mice only. The higher levels of TLR7 in females due to X-chromosome dosage made inhibition of TLR7 activation more difficult. The higher TLR7 expression in female mice significantly increased the viral load but did not lead to eosinophilic disease markers. We hypothesize that this is due to higher type I IFN expression found in females as compared to males in response to respiratory virus infections and to TLR7/8 agonists^54^. Type I IFN expression is known to be protective of IL-5-mediated effects^55^. Overall, these results suggest that TLR7 could be targeted to inhibit RSV infection in males or those with lower *TLR7* expression. As the functions of more TLR7 polymorphisms are identified, their specific detection in the clinic could also serve as a predictor of severe RSV disease, especially in males. Genotyping *TLR7* in patients could be used to determine susceptibility of males to severe RSV disease and asthma development and may inform on therapeutic strategies.

The effects that we observed here are reminiscent of the vaccine mediated enhancement of RSV disease in infants^2^. A recent mRNA vaccine trial reported higher levels of IL-5 in vaccinated infants. If TLR7 is responsible for this enhancement, epigenetically modified oligos or TLR7 inhibitors may help reduce this over reactivity by TLR7. Interestingly, use of TLR7 agonists as vaccine adjuvants enhances RSV replication in mice^56^, and consistently, *Tlr7^−/−^* mice have significantly lower RSV viral loads than WT mice^7^. TLR7 autoimmune disease pathogenesis has been established previously where gain of function of TLR7 lead to the genesis of lupus erythematosus^44^. Our study now shows a role for TLR7 in the pathogenesis of viral infection and disease.

This is not the first report of RSV using a TLR to influence replication. We and others have reported that RSV can interact with TLR4 on target BECs to facilitate RSV entry^4,22,57^. In fact, the RSV-F glycoprotein was the first viral protein identified as a TLR ligand^58^. This suggests that RSV evolved to bind and engage TLR4 because this interaction is likely responsible for the first peak of biphasic ERK MAP kinase activation during RSV entry^22^. The second peak of ERK MAP kinase activation is between 6 and 10 h after inoculation of cells *in vitro*. We speculate that TLR7 activation by RSV replication byproducts likely causes the second later activation of ERK MAP kinase, 8-h after infection, that drives RSV progeny release. More specifically, ERK MAP kinase induction six to eight h after infection causes enhanced RSV-F glycoprotein expression on the plasma membrane and viral replication of infected cells^23^. In summary, RSV has evolved to utilize the kinase signaling that is induced by TLR4 and TLR7 to enhance virus replication and pathogenesis.

TLR7 signaling by RSV could in theory cause expression of type-I interferon and suppression of RSV replication. Negative correlation to RSV viral load in female human NP samples with highest TLR7 expression may be due to an increased influx of professional immune cells with anti-viral effects that overcome the enhancement of RSV infection by TLR7. However, typically early expression of the non-structural proteins (NS) 1 and 2 by RSV effectively suppresses IRF3 and STAT1 signaling by TLR7 and the resulting type-I IFN expression^59^. Consistent with this, our *kika* mouse model shows a type 2 inflammatory response following RSV infection, particularly in males. This phenotype matches what is seen in enhanced RSV disease (ERD), wherein vaccination leads to more severe disease in children^1^. Transcriptional analysis of fatalities in 1967 have found that their immune responses were skewed to a type 2 response.

In summary, our results on TLR7, and the roles of TLR4 in RSV entry shown previously, suggest that RSV has usurped TLR activation and signaling to drive replication. Factors that evolved to protect the host from virus infection have been appropriated by an intracellular pathogen to promote infection of the host. TLR7 signalling is predictive of RSV disease severity and can be targeted with precision antiviral therapies.

## Acknowledgments

Histology tissue processing and staining was performed by Dr. Kacie Norton at the Advanced Microscopy Facility at the University of Alberta. All microscopy images were acquired at the Cell Imaging Core at the University of Alberta. Mouse colony management and mouse work support was provided by the University of Alberta North Campus Animal Service (NCAS) staff. Dr. Kevin Kane provided critical review of the manuscript. HPIV3 was provided by Dr. Ursula Buchholz.

## Ethical Approval

All activities and experimental protocols using human samples and animal experimentation were conducted under the ethical approval and oversight of the human and animal research ethics boards at the University of Alberta; project number: Pro00107301, Pro00099685, and animal project number AUP3799.

## Data Availability

The sequencing data generated in this study will be submitted to the GenBank database in advance of publication or at the request of reviewers.

## Author Contributions

M.J.G, R.D.M., L.M.B., and D.J.M. conceived the study. M.J.G. and D.J.M. wrote the manuscript. M.J.G., R.D.M, L.M.B., and D.J.M designed, performed, and analyzed most experiments. S.H., J.L-O., Q.K., L.D.J, C.D.G, F.E., and D.M.S. performed experiments. C.G.V. provided *kika* mice. O.J. provided support for mass spectrometry. C.D.G., D.M.S., C.G.V., T.C.H., and C.N.J. edited and provided feedback on the manuscript. D.J.M. supervised the study.

## Materials and Methods

### Cells and virus

Bronchial human airway epithelial (1HAEo-) cells were cultured in Minimum Essential Medium (MEM) (GE Healthcare Life Sciences) supplemented with 10% heat-inactivated fetal bovine serum (FBS). HeLa, Madin-Darby canine kidney cells (MDCK), and A549 cells were cultured in Dulbecco’s Modified Eagle Medium (DMEM) (GE Healthcare Life Sciences) supplemented with 10% FBS. During passaging, immortalized cells were detached using 0.25% trypsin with 1mM EDTA. Primary human bronchial epithelial cells (BECs) were obtained from normal, non-transplanted lungs and were cultured in bronchial epithelial growth medium (BEGM; Lonza) supplemented with 5% FBS. At 24 hours prior to experimentation, medium was replaced with BEGM lacking hydrocortisone.

Human RSV strain A2 expressing green fluorescent protein (described previously^60^), was used either directly from infected cell supernatant (media virus) or from virus stocks purified through sucrose gradient ultracentrifugation to remove cellular debris^9^.

HPIV3 was provided by Dr. Ursula Buchholz and is based on the HPIV3 JS strain^61^ and contains an eGFP gene inserted in between the P and M genes. HPIV3 was used directly from infected cell supernatant.

The influenza A virus used was a H1N1 PR8 strain, and was used directly from infected cell supernatant.

### Quantification of RSV and HPIV3 progeny production

1HAEo- cells were seeded in 12 well plates at a density of 1.5×10^5^ cells per well and incubated overnight; primary BEC cells were utilized upon reaching 80% confluency (approximately 10 days post seeding). 1HAEo- or primary BEC cells were infected. At 48 hpi progeny virus was collected and transferred onto HeLa cells. At 4HPI fresh media was replaced onto HeLa cells. At 24HPI HeLa cells were fixed, immunostained, and RSV or HPIV3 infection was quantified as described below.

### Quantification of RSV, HPIV3, and influenza infection

Virus infection was stained and enumerated as described previously^62^. Briefly, at 48 hpi progeny RSV or HPIV3 was collected and transferred onto HeLa cell monolayers, while Influenza A virus progeny was transferred onto MDCK cells. Cells were then fixed and immunostained at 24 hpi. Fixation was performed using methanol:acetone (1:1) for 10 minutes at room temperature. Non-specific antibody binding was blocked via 30-min incubation with PBS supplemented with 5% FBS at room temperature. A polyclonal goat anti-RSV antibody (Meridian), diluted 1:1000 in PBS supplemented with 1% FBS, was selected as the primary antibody used to detect RSV infected cells. A monoclonal antibody targeting human parainfluenza virus type 3 hemagglutinin-neuraminidase protein (Abcam, ab252769), diluted 1:5000 in PBS supplemented with 1% FBS, was selected as the primary antibody used to detect HPIV3 infected cells. A monoclonal antibody targeting influenza A NP (Invitrogen, 7B4G10G8) diluted 1:2000 in PBS supplemented with 1% FBS, was selected as the primary antibody used to detect influenza infected cells. Rabbit anti-goat beta-galactosidase (Abcam, ab136712) was used as a secondary antibody to detect RSV infected cells, and goat anti-mouse beta-galactosidase (Abcam, ab136775) was used to detect HPIV3 and influenza infected cells.

### Drug Treatments

Drug treatment was typically added to cells at the time of infection or at 4HPI, details are found in figure captions. The synthetic oligonucleotide IRS-661 (5′-TGCTTGCAAGCTTGCAAGCA-3′) with phosphorothioate backbone; Integrated DNA Technologies), was described previously to antagonize TLR7 stimulation^63^. ODN-SCRAM is made from the scrambled sequence of IRS-661 (5′-GCTCCTAGTAGAGATCTCAG-3′)^64^. UO126 (Tocris), was used as a MEK inhibitor for downstream ERK inhibition. SCH772984 (Cedarlane) was used as a direct ERK inhibitor. Loxoribine (Sigma) is a guanosine analogue TLR7 agonist. TJ-M2010-5 (MedChemExpress) was used as a MyD88 inhibitor. Nucleoside analogues 5-fluorocytidine (5-FC), bromodeoxyuridine (BrdU), 2-iodoadenosine (2-IA), N4-benzyolcytidine (N4BC), and ribavirin were purchased from Sigma-Aldrich. CL097 (Sigma) was used as a dual TLR7 and 8 agonist.

### Antibodies

RSV-G (polyclonal goat anti-RSV antibody (Meridian, B65860G-1)), SOD1 ((Santa-Cruz, FL-154), P^S422^ eIF4B (CST, 3591), P^S112^ BAD (CST, 9291), ERK (CST, 4695), P-ERK (CST, 4370), p90 RSK (CST, 9355), and P^S380^ p90 RSK (CST, 11989) were detected by western blot. TLR7 was detected by western blot using a N-terminal monoclonal antibody (CST, 5632S) and a C terminal polyclonal antibody (Invitrogen PA1-28109). TLR7 was detected for histology and confocal imaging using a C-terminal polyclonal antibody (Invitrogen, PA1-28109). RSV-F, N, and P were detected for confocal imaging using an anti-RSV3 monoclonal antibody (Leica). Endosomal markers were detected for confocal imaging using the following antibodies: LAMP1 (CST, 9091), LAMP2 (Abcam, EPR19512), Rab5c (CST, 3547), Rab7 (CST, 9367), and LC3 (CST, 12741). Lysotracker (Thermo Fisher Scientific) was used for confocal microscopy to stain acidic compartments, and DAPI (Abcam, ab228549) was used to stain DNA.

### Base modified RNA Additions

The first 20 nucleotides of the RSV-N gene were synthesized as an RNA fragment (AUGGGCAGCAAUUCAUUGAG) (Integrated DNA Technologies) as a control. The same sequence was also synthesized as an RNA fragment with N^6^-methyladenosine modifications (AUGGGC/iN6Me-A/GC/iN6Me-A//iN6Me-A/UUC/iN6Me-A/UUG/iN6Me-A/G) (Integrated DNA Technologies). Both RNA fragments were added in a concentration gradient to 1HAEo-cells at 4HPI, and at 48HPI progeny virion release was quantified as described above.

### Quantification of RSV via RT-qPCR

Briefly, RNA was extracted using a Nucleospin RNA plus kit (Macherey-Nagel, 740984.50). The reaction was prepared using qScript™ XLT One-Step RT-qPCR ToughMix (QuantaBio, CA75994-402) with primers designed to RSV^65^ and housekeeping gene hypoxanthine guanine phosphoribosyltransferase (HPRT) (Life Technologies, Hs02800695_m1). The reaction was run on the 7500Fast Real-Time PCR System (Applied Biosystems). We have previously described the determination of equivalent FFU/mL of RSV from Ct values based on comparison to a standard curve^43^. For other experiments, RSV expression was determined relative to HPRT by calculating 2^-ddCt^.

### Fluorescence microscopy

Our techniques for fluorescence microscopy have been described previously^66^. A spinning disc confocal microscope (Wave FX #2, Quorum Technologies) running Volocity software (PerkinElmer) in the Cell Imaging Centre at the University of Alberta was used for acquisition.

### Cell lysate collection and western blot

Mos LB, containing Mini EDTA-free Protease Inhibitor cOmplete cocktail (Roche), was added to cells and incubated on ice for 20 minutes. Protein concentration in cell lysates were measured by Qubit (Life Tech) according to the manufacturer’s protocol, and lysates were stored at -80°C. 4-15% pre-cast polyacrylamide gels were loaded with five parts lysate added to 1 part 6x Laemmli Sample Buffer with 5% β-mercaptoethanol that were boiled for 10 minutes at 100°C. Protein was transferred to nitrocellulose membrane (PALL Life Science), blocked in TBS-T plus 5% bovine serum albumin (BSA; Sigma), and incubated with primary antibody for 1 hour at room temperature or overnight at 4°C. Primary antibodies were detected via a horseradish peroxidase conjugated secondary antibodies. Chemiluminescent Pierce ECL substrate (Fisher) was used to detect protein and blots were imaged on an Chemidoc imager (Bio-Rad).

### siRNA knockdown of TLR7

1HAEo-cells were seeded in 12 well plates at a density of 4×10^4^ cells per well and incubated for 48 hours prior to transfection. Immediately prior to transfection, 1HAEo-cells were washed with PBS and exactly 1mL of fresh media was replaced onto cells. Following the manufacturer’s protocol, 1HAEo-cells were transfected at a confluency of approximately 50% with siRNA at a final concentration of 15 nM. Briefly, AllStar Negative Control siRNA (proprietary target sequence; Qiagen), AllStar Hs Cell Death siRNA (proprietary mixture of oligonucleotides; Qiagen), or TLR7 (Hs_TLR7_6 target sequence: 5’-CAGACCTTGGATCTAAGTAAA-3’; Qiagen) was added to 100 µL of jetPRIME buffer (Polyplus) and vortexed briefly. 3 µL of jetPRIME reagent was added to each reaction mix; transfection mixes were vortexed briefly and incubated at room temperature for 10 minutes. Transfection mixtures were added to respective wells in a dropwise manner. Following transfection, 1HAEo-cells were incubated for 48 hours prior to experimentation to enable knockdown of target protein and recovery of cells from transfection protocol. Transfection efficiency was estimated by examining cell rounding and overall monolayer damage in wells transfected with AllStar Cell Death siRNA. Cell rounding and monolayer damage was imaged via light microscopy with a 10x objective lens using the EVOS FL Auto (Life Technologies) microscope.

### Mass Spectrometry

Primary BECs were grown in a T75 flask to confluency, then washed with PBS and lifted using 0.25% trypsin-EDTA and resuspended in growth media. Resuspended cells were then transferred to a 15mL conical tube (Falcon) and centrifuged at 800x g for 10 min to pellet the cells. Media containing trypsin was aspirated and cells were resuspended in PBS before centrifuging at 800x g for 10 min. PBS was then aspirated, and the cell pellets were stored at -80°C.

To help enrich the cell lysates for membrane proteins, samples were processed with S-Trap Mini Columns (Protifi, #C02-mini-10). This involved solubilization in 5% SDS, acidification to pH < 1, and exposure to high concentration methanol. Trypsin was then used to digest the proteins bound to the S-Trap column and the peptides are eluted. Digested peptides were run on an Orbitrap Fusion Lumos Tribrid Mass Spectrometer (Thermo Fisher Scientific) in the Alberta Proteomics and Mass Spectrometry Facility at the University of Alberta.

### Nanostring nCounter

Primary BECs from two different donors were seeded in BEGM (Lonza, #CC-3170) supplemented initially with 5% FBS in 12-well plates and incubated until they reached approximately 80% confluency. Cells were then infected with sucrose-purified RSV at an MOI of 1 in BEGM. At the time of infection, cells were treated with 1µM IRS-661 or left untreated. At 48 HPI, RNA was harvested separately from all wells with a NucleoSpin RNA Plus Kit (Macherey-Nagel, MN 740984.250). 100ng of high-purity RNA per sample was analyzed with the NanoString nCounter Host Response to assess gene expression (Bruker Spatial Biology, #100052).

### RNA base modification sequencing

Primary human BECs were seeded in 6 well plates, infected with RSV at an MOI of 0.5, and treated with DMSO or 10μM U0126 (Tocris, 1144/5) at 4 hours post infection. At 48 hours post infection, cells were harvested and RNA extracted using a Nucleospin RNA plus kit (Macherey-Nagel, 740984.50). Ribosomal RNA was then removed using NEBNext rRNA Depletion Kit v2 (New England Biolabs, E7400L). Sequencing libraries were prepared using the Direct RNA Sequencing Kit (Oxford Nanopore Technologies, SQK-RNA004). Libraries were sequenced on a MinION flow cell (RNA) (Oxford Nanopore Technologies, FLO-MIN004RA) using the MinION Mk1B sequencer. Base calling was performed using MinKNOW software set to super accurate (https://nanoporetech.com/document/experiment-companion-minknow#quick-start-guides-introduction). Passed reads had a minimum quality score of 10. Modified base calling was set to call inosine, m6A, m5C, and pseudouridine in all contexts. Passed bam files were aligned to the RSVA2 genome (GenBank: KT992094.1, ^67^) using Dorado (https://github.com/nanoporetech/dorado). Modkit was used to identify base modifications in these aligned files (https://github.com/nanoporetech/modkit).

### TLR7 Inhibitor Treatments In Vivo

4- to 6-week-old BALB/c mice (The Jackson Laboratory) were anesthetized with isoflurane and 20μL of inoculum was administered intranasally. The inoculum contained: 1×10^6^ FFU sucrose-purified RSV-GFP or 35% sucrose (mock-infection), 15 nmol IRS-661 or 15 nmol ODN-SCRAM control. Mice throats and nares were swabbed daily with PurFlock Ultra Flocked Swabs (Puritan, #89194-898) which were placed in 400 µL PBS and then harvested with a NucleoSpin Virus Kit (Macherey-Nagel, #740983.250). 15 nmol of IRS-661 or ODN-SCRAM in PBS were re-administered daily under isoflurane anesthesia. Mice were euthanized with CO_2_ at 3 or 5 days post-infection, and lungs were dissected. The superior, middle, and inferior lobes were harvested and fixed in 1mL of 36.5-38% Formaldehyde solution (Sigma Aldrich, F8775) for histological processing and analysis. The left lung and post-caval lobes were harvested, homogenized, and spun at 4303xg for 10min. RNA was extracted from the lung supernatant using a NucleoSpin Virus Kit (Macherey-Nagel, MN-740983.250). RNA was analyzed by RT-qPCR to quantify viral loads, and viral copy number was determined in comparison to a standard curve as described previously^43^. RSV copy number was then normalized to the input RNA in nanograms (ng). RNA was also used to determine lung cytokine gene expression using RT-qPCR as described above.

### Kika Mice Infections

A C57Bl/6 TLR7^Y264H^ (*kika*) mouse breeding pair were obtained as a gift from Dr. Carola G. Vinuesa (Francis Crick Institute, United Kingdom). Mice were bred by staff at the University of Alberta North Campus Animal Service (NCAS). 4–5-week-old mice were infected with 20μL of 1×10^6^ FFU RSV media by intranasal inoculation under isoflurane anesthesia. Mice were then monitored and weighed daily for 72 hours. At 72 HPI, mice underwent euthanasia by CO_2_ inhalation. Blood within the thoracic cavity was harvested and into tubes containing 30μL EDTA, 0.5M pH 8.0. Lungs were dissected and the superior, middle, and inferior lobes were harvested and fixed in 1mL of 36.5-38% Formaldehyde solution (Sigma Aldrich, F8775) for histological processing and analysis. Blood was spun at 2,000xg to isolate plasma for cytokine analysis. The left lung and post-caval lobes were flash-frozen in liquid nitrogen and homogenized with a pestle prior to RNA extraction using a NucleoSpin RNA Plus Kit (Macherey-Nagel MN740984.250). RSV expression was determined for each lung RNA sample using RT-qPCR as described above and normalized to housekeeping gene hypoxanthine guanine phosphoribosyltransferase (HPRT) by calculating 2^-ddCt^. Mice lacking a detectable viral load were excluded from analysis. Lung RNA was also used for bulk RNA sequencing.

### Plasma cytokine multiplex quantification

Plasma was analyzed using the ProcartaPlex™ Mouse Immune Monitoring Panel, 48plex (EPX480-20834-901). Results were quantified using a Luminex MAGPIX system. Cytokine concentrations are reported as net median fluorescent intensity (MFI)^68^.

### Histology

Fixed lung sections were embedded in paraffin, sectioned serially with a microtome, and then mounted on slides. Prior to staining, samples were deparaffinized and rehydrated. Samples were incubated in 0.01M citrate buffer (pH 6) for antigen retrieval. A 30% H_2_O_2_ wash was performed to inhibit endogenous peroxidase activity before blocking samples in PBS-T containing 3% BSA. The mounted lung sections were then stained with anti-TLR7 (1:250) (Invitrogen, PA1-28109). and donkey anti-rabbit HRP light-chain (1:200) antibodies and processed with a DAB Substrate Kit (Thermo Fisher Scientific PI34002). Hematoxylin was used as a counterstain. Tissue processing and staining was performed by Dr. Kacie Norton at the Advanced Microscopy Facility at the University of Alberta. Histology slides were then imaged with a Zeiss Axioscan.Z1 Slide Scanner at 40X magnification in the Cell Imaging Centre at the University of Alberta. TLR7 staining was quantified using QuPath^69^. As a control, recombinant TLR7 protein (Cedarlane 9567-TR-050) was incubated with TLR7 antibody (Invitrogen PA1-28109) at a ratio of 5:1. Protein-saturated TLR7 antibody was then used to stain mounted lung samples. Secondary antibody staining, DAB-substrate processing, and hematoxylin counter-staining were performed normally, as described above.

### Lung Bulk-RNA sequencing

Mouse lungs were harvested at 3 days post infection and total RNA was extracted using a Nucleospin RNA plus kit (Macherey-Nagel, 740984.50). Total RNA was then sent to Genewiz From Azenta Life Sciences for polyA selection bulk RNA sequencing with 30 million paired end reads per sample on an Illumina platform. Read quality control was performed using FastQC (Andrews, S. (2010). FastQC: A Quality Control Tool for High Throughput Sequence Data [Online]. Available online at: http://www.bioinformatics.babraham.ac.uk/projects/fastqc/). Trimming was performed using Fastp^70^. Reads were aligned to the GRCm39 reference genome (Genome Reference Consortium. (2020). *Mus musculus (house mouse) genome assembly GRCm39* (NCBI RefSeq assembly accession GCF_000001635.27). National Center for Biotechnology Information. Retrieved from https://www.ncbi.nlm.nih.gov/datasets/genome/GCF_000001635.27/) and genes were also counted using STAR^71^. DESeq2 was used to analyze differential gene expression^72^.

### AlphaFold 3 TLR7 modelling

The reference gene sequence for human TLR7 (Gene ID: 51284) was accessed from the RefSeq database (National Library of Medicine). AlphaFold 3 (Google DeepMind) was then used to predict and model the protein structure of TLR7^wt^ homodimers.

### Analysis of TLR7 RNA expression by single cell sequencing in human lungs

The results here are in whole or part based upon data generated by the LungMAP Consortium and downloaded from (www.lungmap.net) on May 27, 2025. Single-cell transcriptomic signatures of 46500 normal human lung cells from donors aged 1 day to 33 years old were originally generated by ^17^; https://doi.org/10.7554/eLife.62522; Dataset ID: LMEX0000004388). Further independent analysis of this data set was completed using LungMAP ShinyCell software.

